# Chromosome-level genome assemblies for the latent pine pathogen, *Diplodia sapinea*, reveal two rapidly evolving accessory chromosomes

**DOI:** 10.1101/2025.03.26.645563

**Authors:** Preston L. Shaw, Bernard Slippers, Brenda D. Wingfield, Benoit Laurent, Benjamin Penaud, Michael J. Wingfield, Pedro W. Crous, Wubetu Bihon, Tuan A. Duong

## Abstract

*Diplodia sapinea* (Dothideomycetes) is a latent fungal pathogen with a global distribution that predominantly infects *Pinus* species. The impact of the fungus is increasing due to climate-driven range expansion and thus wide-scale disease outbreaks. With the aim of developing high quality genome resources, we generated chromosome-level genome assemblies for three *D. sapinea* isolates and low-coverage Illumina genome data for six additional isolates. By comparing these genome assemblies, we identified 14 core chromosomes and two accessory chromosomes (ACs) in the pathogen. The ACs encode 80 and 155 proteins, respectively, while 11374 - 11609 genes were identified in the core chromosomes. Both ACs had lower gene density and higher proportions of transposable elements compared to the core chromosomes. Sequence analysis indicated that genes on the ACs are rapidly evolving, suggesting they serve as evolutionary hotspots in the species. Sequence homology analyses suggested that the ACs were likely acquired horizontally, probably from a species in the Dothideomycetes. We designed PCR-based assays to aid in the detection of the ACs and applied these on a set of 37 isolates from 14 countries. One of the ACs was detected in 33 isolates from 13 countries, while the other AC was absent in all isolates tested. Pathogenicity trials on *Pinus patula* seedlings showed no correlation between the presence of ACs and isolate aggressiveness. The high-quality genomes provided here offer important resources for future research on this globally important pathogen, including the biological roles of the ACs.

## Introduction

*Diplodia sapinea* (synonyms = *Diplodia pinea*; *Sphaeropsis sapinea*) is an opportunistic ascomycete fungal pathogen of conifers that predominantly infects *Pinus* spp. (Luchi et al. 2014). The pathogen has a global distribution (Wingfield et al. 2025) and is thought to have spread to areas where non-native *Pinus* spp. are planted through the trade of infected pine germplasm (Burgess and Wingfield 2002; Wingfield et al. 2001). In recent years, damage due to *D. sapinea* has increased in areas not previously affected including those where it was previously not known to occur, such as northern parts of Europe, likely due to climate change (Brodde et al. 2019; Müller et al. 2019).

*Diplodia sapinea* can persist in asymptomatic trees for extended periods of time, potentially indefinitely, prior to disease development (Bihon et al. 2011; Stanosz et al. 2001). Disease symptoms develop in trees that become physiologically stressed or damaged due to damage from hail, drought stress or other unfavourable environmental conditions, as well as biotic disturbances (Palmer et al. 1987; Stanosz et al. 2001; Zwolinski et al. 1990). Symptoms associated with *D. sapinea* infection include shoot dieback, stem cankers, resinosis, blue staining and root disease, and in severe cases, these can result in host mortality (Chou 1976; Swart et al. 1987; Wingfield and Knox-Davies 1980). Severe disease outbreaks can cause extensive damage to plantations of *Pinus* spp. by killing new shoots, causing deformation and decreasing wood quality (Brodde et al. 2019; Caballol et al. 2022; Swart and Wingfield 1991; Zwolinski et al. 1990).

There are currently five genome assemblies available for *D. sapinea* in the NCBI genome database (Bihon et al. 2014; van der Nest et al. 2014; Yu et al. 2022). The assemblies are highly fragmented, with the number of scaffolds ranging from 931 to 5957 and assembly N50 ranging from 39,4 to 131,4 Kb. Yet the availability of these genome resources has enabled studies on the biology and genetics of the species. The first of these studies characterised the mating type genes and was used in the development of mating type diagnostic markers (Bihon et al. 2014). By comparing the genome assemblies of two isolates, the authors showed that *D. sapinea* is heterothallic, characterised by the presence of either *MAT1-1* or *MAT1-2* idiomorph in each of the isolates, and the novel *MAT1-2-5* gene was found. Bihon et al. (2014) also developed mating type markers and used these to examine the mating type distribution in natural populations of the fungus, suggesting the occurrence of a cryptic sexual cycle in *D. sapinea*. The genome resources of *D. sapinea* have also been used to develop a transformation protocol allowing for reverse genetic studies (Oostlander et al. 2024).

Many species of fungi have compartmentalised genomes consisting of core and accessory genomes (Covert 1998; Galazka and Freitag 2014). The accessory genome can include accessory regions on core chromosomes (CCs) and/or ACs. Accessory chromosomes, also referred to as ‘B’ chromosomes, supernumerary chromosomes or accessory chromosomes, are additional chromosomes that are not present in all individuals of a given species (Bertazzoni et al. 2018). Accessory chromosomes are thought to arise via horizontal chromosome transfer and/or vertically through duplication events of the core genome or a combination of both processes (Soyer et al. 2018). This form of genome compartmentalisation provides the advantage that essential processes can be maintained by the core genome while the accessory genome can rapidly evolve and potentially acquire new beneficial functions (Croll and McDonald 2012). This is thought to be particularly true and of relevance in plant pathogen evolution, and therefore, an important question to also consider in *D. sapinea* where host-pathogen interactions need to be better understood.

As in other plant pathogens, the growing availability of genomes in the *Botryosphaeriaceae* in the last decade has led to significant new insights regarding the biology and host-pathogen interactions, in these fungi (Belair et al. 2023; Nagel et al. 2021a, b; Yu et al. 2022). Nagel et al. (2021b) used these genomes to map the evolution of the reproductive strategy in the *Botryosphaeriaceae*, showing multiple independent transitions between heterothallism and homothallism. Comparative genomics studies amongst species of the *Botryosphaeriaceae* and other Dothideomycetes have also revealed an abundance (amongst the highest in the Class) of secreted hydrolytic enzymes and secondary metabolite BGCs in the genomes of *Botryosphaeria*, *Macrophomina*, *Lasiodiplodia*, and *Neofusicoccum*, showing similarity to genomic profiles to other necrotrophic plant pathogens (Nagel et al. 2021a, Yu et al. 2022). Interestingly *Diplodia* species, including *D. sapinea*, had the lowest numbers of pathogenicity- and virulence-related genes. Nagel et al. (2021b) found that the genomes in the *Botryosphaeriaceae* are not compartmentalised in terms of genome evolution, as is seen in some other plant pathogens. None of these *Botryosphaeriaceae* genomes, however, were assembled to chromosome level, and the potential influence of accessory chromosomes (ACs), which are known to play an important part in plant pathogen evolution (Witte et al. 2021) could not be considered.

In this study, we generated three chromosome-level genome assemblies of *D. sapinea* and used these as a basis to identify core and potential accessory chromosomes. The composition, structure and evolution of the chromosomes were investigated. We also developed molecular markers to characterise the presence of the identified ACs and considered their possible role in the pathogenicity of the species.

## Materials and methods

### Cultures

Isolates of *D. sapinea* used in this study (Supp. Table S1) were sourced from the culture collection (CMW) of the Forestry and Agricultural Biotechnology Institute (FABI) at the University of Pretoria, South Africa and the CBS-KNAW Fungal Biodiversity Centre, Westerdijk Fungal Biodiversity Institute, Netherlands. Single hyphal tip cultures were made for all the isolates and resulting cultures were maintained on MEA at 25 °C.

### Nanopore and Illumina genome sequencing

Three isolates of *D. sapinea* (CMW190, CMW39103, CMW45410) were selected for Oxford Nanopore long-read sequencing on the MinION device. These isolates were grown in YM broth (2% malt extract, 0,5 % yeast extract) for 3 – 5 days until sufficient mycelial mass was obtained. The mycelium was freeze-dried and ground into a fine powder in liquid nitrogen. DNA was extracted from ground mycelium using QIAGEN Genomic-tips (QIAGEN, Hilden, Germany). The cells were lysed for 4 hours at 42 °C in cell lysis buffer (10 mM Tris-HCl pH 7.9; 20 mM EDTA, 1 % Triton® X-100, 500 mM Guanidine-HCl, 200 mM NaCl) supplemented with 0,5 mg/ml cell lyzing enzyme from *Trichoderma harzianum* (Sigma-Aldrich, Missouri, USA), 0,8 mg/ml proteinase K (Sigma) and 20 ug/ml RNase A (QIAGEN). The DNA was then purified from the cell lysate using the QIAGEN G-20 Genomic-tip following the manufacturer’s suggested protocols.

For long-read sequencing, a genomic library was prepared for each isolate using the Genomic DNA by Ligation (SQK-LSK109) library preparation kit (Oxford Nanopore Technologies, Oxford, UK) and sequenced on an R9.4.1 flow cell using the MinION sequencing platform (Oxford Nanopore Technologies). Base calling was performed with Guppy v2.7.3 (https://community.nanoporetech.com) using the flip-flop model. Illumina sequencing was conducted on the Illumina HiSeq platform. A TruSeq PCR-free library was constructed for each isolate and sequenced to obtain 151 bp paired-end reads.

### Genome assembly and identification of accessory chromosomes

Nanopore data were trimmed using Porechop v0.2.4 (https://github.com/rrwick/Porechop) to remove any remaining sequencing adapters. Two draft genomes were generated for each isolate using Canu v.2.0 (Koren et al. 2017) and Flye v2.3.1 (Kolmogorov et al. 2019). The draft genome assemblies for each isolate were then aligned to one another using MUMmer v3.23 (Kurtz et al. 2004) to confirm collinearity and identify any conflicts (i.e. large structural variations) between them. Identified conflicts were resolved by aligning the Nanopore data to both draft assemblies using Burrows-Wheeler Aligner (BWA) v0.7.17-r1188 (Li and Durbin 2010) and manually curating the regions with inconsistencies between the assemblies. The alignment files were converted to BAM format using samtools v1.13 (Li et al. 2009) and viewed using Integrative Genomics Viewer (IGV) v2.8 (Robinson et al. 2011). A final gap-filling step of the curated genome assemblies was done using PBJelly 15.8.24 (English et al. 2012).

The final curated long-read assembly for each isolate was polished using Illumina sequence data. Quality and adapter trimming of the Illumina sequence data was done using Trimmomatic v0.33 (Bolger et al. 2014). Each assembly was initially polished with three iterations using Pilon v1.23 (Walker et al. 2014), followed by a final round of polishing using Racon v1.4.12 (Vaser et al. 2017).

The completeness of the final assemblies was determined with BUSCO v4.0.5 (Simão et al. 2015) using the dothideomycetes_odb10 lineage dataset. To determine whether any of the assembled contigs represented complete chromosomes, the sequences were scanned for the telomeric repeat motif (CCCTAA) using Bowtie v1.3.1 (Langmead et al. 2009). Only sequences containing at least 10 consecutive repeat motifs were classified as a telomeric region. Accessory chromosomes were identified by aligning each complete genome assembly to each other using MUMmer v3.23 (Kurtz et al. 2004). Their presence was further confirmed by mapping short reads from isolates lacking the ACs onto the reference genome that contained them. The mapped reads were then inspected in IGV (Robinson et al. 2011) to assess the presence or absence of coverage over the AC regions.

### Genome annotation

Soft-masking of the genomes was performed using RepeatMasker v4.0.7 with a repeat library identified with the REPET package (Flutre et al. 2011). Annotation of protein-coding genes was generated with BRAKER v2.1.5 (Hoff et al. 2019) using the fungi_odb10 protein database with additional proteins from *M. phaseolina*, *D. corticola*, *D. seriata* and *N. parvum* downloaded from GenBank as evidence for gene prediction. Initial protein hints were generated using ProtHint v2.5.0 (https://github.com/gatech-genemark/ProtHint) prior to running BRAKER. Functional annotation of the predicted proteins was done using the Blast2GO (Conesa et al. 2005) plugin as part of the CLC Genomics Workbench program v22.0.2. The Blast2GO results were used for comparing the proteins encoded by the CCs and ACs with regard to similarity scores and the top hit species in the NR protein database. To test for functional enrichment between genes on the ACs and the core chromosomes, gene ontology (GO) enrichment analysis was performed with Blast2GO using the Fischer’s Exact test with default parameters.

### Genomic variant discovery in ACs

To determine the prevalence of the ACs and the genetic variation in AC genomic sequences, low coverage Illumina sequence data for six additional *D. sapinea* isolates from five countries, as well as the three originally sequenced isolates (Supp. Table S1), were aligned to the reference assembly of isolate CMW45410 using Burrows-Wheeler Aligner (BWA) v0.7.17-r1188 (Li and Durbin 2010). The alignment files were converted to BAM format using samtools v1.12. The alignment files were then processed using freebayes v1.3.8 (Garrison and Marth 2012) to identify SNPs throughout the genome of *D. sapinea*. Bcftools (Danecek et al. 2021) was used to filter identified SNPs for quality control concerning depth (DP < 3) and probability score (GQ > 20). Variants were intersected with annotated genomic features using bedtools v2.26.0 (Quinlan and Hall 2010). The SNP density was then calculated for CDS regions and compared between genes on the ACs, a random subset of 80 genes from core chromosomes and 80 random BUSCO genes.

### Secondary metabolite gene cluster, effector predictions

The generated annotations were used as support for secondary metabolite gene cluster predictions with the fungal version of antiSMASH v7.0.0 (Blin et al. 2023) using a relaxed strictness and the CASSIS feature turned on. Signal peptides were predicted from the putative amino acid sequences using SignalP v5.0 (Almagro Armenteros et al. 2019). Unconventionally secreted peptides were identified using OutCyte (Zhao et al. 2019). The predicted secretome, as identified by both SignalP and OutCyte, was subject to effector prediction using EffectorP-fungi 3.0 (Sperschneider and Dodds 2022).

### Orthology analysis

Protein clustering was done with OrthoFinder v2.5.4 (Emms and Kelly 2019) using protein sequence data from three *D. sapinea* isolates (CMW39103, CMW190 and CMW45410) for which whole genome and annotation were generated, as well as using proteomes from additional species of the *Botryosphaeriaceae* for which data was publicly available at the time of this study (Supp. Table S2). OrthoFinder analyses were run with a range of MCL inflation parameters (from 1.5 to 4.0 using increments of 0.5), and the run that produced the highest number of single-copy orthogroups was selected for further analyses. The orthogroups generated by OrthoFinder were used to identify and compare shared and unique genes on the ACs and core chromosomes.

### Phylogenomic analysis of *Botryosphaeriaceae*

The genomes of all *Botryosphaeriaceae* species used for protein clustering were analysed with BUSCO using the dothideomycetes_odb10 lineage data set. Single-copy BUSCO genes that were shared amongst all 30 species were identified and used to construct a species phylogeny using a coalescence approach. Shared, single-copy BUSCO amino acid sequences were extracted from the BUSCO outputs and aligned using PRANK v.170427 (http://wasabiapp.org/software/prank/) with default parameters. The aligned datasets were trimmed with trimAI v1.5 (Capella-Gutiérrez et al. 2009) with the “-atomated1” option. Maximum Likelihood (ML) trees were inferred for individual datasets using IQ-TREE v1.6.11 (Nguyen et al. 2015) with best fit substitution models computed automatically and 1000 rapid bootstrap replicates. The resulting ML trees were used to construct a coalescence tree using ASTRAL v5.7.7 (Zhang et al. 2018). The species tree obtained from ASTRAL was optimised for branch length with RaxML v8.2.11 (Stamatakis 2014) using a concatenated dataset derived from aligned and trimmed individual BUSCO datasets. Gene concordance factors (gCF) were computed in IQ-TREE using the individual gene trees.

### Development of diagnostic multiplex PCR assays for the presence of ACs

A total of 7 primer pairs were designed for multiplex PCR assays (Supp. Table S3). Genes used for primer design to detect AC Chr 15 were chosen based on their sequence conservation in isolate CMW190 as well as their position on the chromosome. This was done by comparing Illumina sequence alignments of the additional *D. sapinea* isolates (Supp. Table S1) to AC Chr 15 in IGV. As AC Chr 16 was unique to one isolate, the method of using conserved sequences was not applicable. Gene candidates for primer design were thus chosen based on whether they formed single-copy orthogroups after protein clustering with OrthoFinder. Three primer pairs were designed to amplify gene regions on AC Chr 15. These regions included a gene encoding a secreted hypothetical protein, a polyketide synthesis gene, and a toxin synthesis gene. Furthermore, three primer pairs were designed to amplify gene regions on AC Chr 16, targeting genes encoding a putative C6 finger domain protein, a hypothetical protein, and a NACHT and WD40 domain. The final primer pair was designed to amplify the beta-tubulin gene regions, acting as a positive control. All primers were designed using Primer3Plus v2.4.2 (Untergasser et al. 2012) to have similar annealing temperatures and different amplicon sizes to allow PCR multiplexing (Supp. Figure 1).

### PCR amplification of Accessory Chromosome

A total of 37 isolates from 14 countries across 6 continents were used in PCR assays to detect the presence of the ACs (Supp. Table S4). Three isolates (CMW39103, CMW190 and CMW45410) for which chromosome-level genome sequences had been generated were included as controls. DNA samples were extracted from pure cultures actively growing on MEA (2% malt extract, 2% Agar, Biolab, South Africa). Mycelium was transferred to separate 1.5 ml Eppendorf tubes, and DNA was extracted using PrepMan Ultra Sample Preparation Reagent (Thermo Fisher Scientific) using a temperature of 95°C for a total of 10 mins. The mycelium was macerated using a pestle after the first 5 mins of heating. The extracted DNA was then diluted five times with 10 mM Tris-HCl, pH 8.0, and subsequently used for PCR amplification. A multiplex PCR assay was performed for the detection of each AC separately. Each PCR reaction consisted of 2 µl DNA template, 0.2 µl FastStart Taq DNA polymerase (Roche Applied Science, Mannheim, Germany), 2.5 µl dNTPs (2 mM each), 2.5 µl 10x PCR buffer w/ MgCl2, 5 µl GC-Solution, 0.5 µl MgCl2 (25mM), 0.5 µl of each primer (10 µM) and H_2_O up to the final volume of 25 µl. PCR amplification was carried out with a MyCycler Thermal Cycler from Bio-Rad using cycling conditions as described by de Beer et al. (2014) with an annealing temperature of 55°C. The final PCR products were stained with GelRed (Biotium, California, United States) and separated using gel electrophoresis in a 2% agarose gel (SeaKem LE Agarose, Lonza Bioscience) at 80 V for 45 minutes. The PCR amplicon was then visualised under UV light using a Gel Doc EZ Imager (Bio-Rad). The presence/absence of the ACs was identified by the presence/absence of the amplicons from the targeted genes. The identity of each isolate was confirmed by sequencing the positive control amplicon targeting the beta-tubulin gene region and conducting a BLAST analysis against the NCBI non-redundant nucleotide database.

### Pathogenicity trial

A pathogenicity test was conducted on *Pinus patula* saplings, approximately two-years-old using six isolates of *D. sapinea* (Supp. Table S5). Isolates used in the pathogenicity trial were chosen based on whether they possessed the ACs or not, as indicated by the PCR screening. Isolates used in the pathogenicity trials were freshly prepared on 2% MEA and maintained at room temperature until the day of inoculation. *Pinus patula* saplings were inoculated and maintained in a greenhouse d at 25 °C.

Inoculations were performed by removing the outer bark to expose the cambium of the plants with a 4 mm diameter cork borer approximately 5 cm below the apical bud. A plug of mycelium-covered media was placed onto the freshly made wound, mycelium side facing the wounded surface, and sealed with parafilm to reduce desiccation and contamination. A total of seven saplings were inoculated per isolate, and an equal number were inoculated with 2% MEA as a negative control. The plants were then left for a period of three weeks. The resulting lesions were measured by cutting away the bark with a scalpel and measuring the length of the lesions downwards on the stems starting from the base of the inoculation point. Re-isolations were made from the lesions for three plants per isolate by placing a small piece of stem tissue harvested from the base of the lesion on 2% MEA. Fungi that resembled *D. sapinea* in culture were transferred to fresh growth medium, purified and identified via PCR amplification and sequencing of the ITS region using the ITS-1F and ITS-4 primers.

## Results

### Chromosome level assemblies reveal the presence of two accessory chromosomes in *D. sapinea*

The long reads generated for three isolates of *D. sapinea* (CMW39103, CMW190 and CMW45410) were used to assemble the genomes using two different assemblers. In most cases, Canu generated the best initial assembly results but slightly under assembled the genome, whereas Flye performed well but occasionally mis joined contigs (Supp. Table S6). The Canu assemblies were selected as the base assemblies for the curation process by means of whole genome alignment and long read-mapping for manual verification (Supp. Figure S2). The final genome assemblies of three *D. sapinea* isolates had 14, 15 and 16 contigs, respectively, with assembled genome sizes ranging from 37.01 to 38.4 Mb (Table. 1). The telomeric repeat sequence (CCCTAA) was detected at both ends of 15 contigs in at least one of the three assembled genomes (Supp. Figure S3) indicating that these genomes were likely assembled into complete or close to complete chromosomes. However, no telomeric regions were detected for the Chr 16 in CMW45410. BUSCO analyses using the dothideomycetes_odb10 dataset (n=3786) indicated a high level of assembly completeness in all three assemblies with BUSCO scores higher than 99% (Table 1). Genome annotation resulted in a total of 11485 protein-coding genes in CMW39103, 11770 in CMW190 and 11609 in CMW45410. Of the predicted protein coding genes, 79 are located on Chr 15 in CMW190, 80 on Chr 15 in CMW45410, and 155 on Chr 16. BUSCO completeness for the genome annotations using the dothideomycetes_odb10 dataset were 98.3%, 98.4% and 97.8%, respectively.

**Table 1.** Assembly and annotation statistics of CMW39103, CMW190 and CMW45410 generated in this study.

|  | CMW39103 | CMW190 | CMW45410 |
| --- | --- | --- | --- |
| Genome size (bp) | 37011478 | 38053143 | 38401314 |
| GC content (%) | 56.66 | 56.57 | 56.54 |
| Number of contigs | 14 | 15 | 16 |
| N50 (bp) | 2852041 | 2851318 | 2860161 |
| L50 | 6 | 6 | 6 |
| BUSCO (Dothideomycetes: n:3786) | 99.4% | 99.4% | 99.3% |
| Accessory chromosomes and sizes | - | Chr 15 (0.48 Mb) | Chr 16 (0.64Mb);<br>Chr 15 (0.46Mb) |
| No. protein coding genes | 11485 | 11770 | 11609 |
| No. single exon genes | 2333 | 2414 | 2374 |
| Mean CDS length (b) | 1463 | 1456 | 1458 |
| Mean exons per gene | 3.0 | 3.0 | 3.0 |
| Mean exon length (b) | 487 | 487 | 488 |
| Mean intron length (b) | 85 | 85 | 87 |

Synteny comparison between three assembled genomes revealed a high level of collinearity between three genomes, with end-to-end alignment on 14 out of the 16 pseudo-chromosomes (Figure 1). An additional pseudo-chromosome (Chr15) was shared by CMW190 and CMW 45410, and another (Chr16) was unique to CMW45410 (Figure 1).

**Figure 1.**
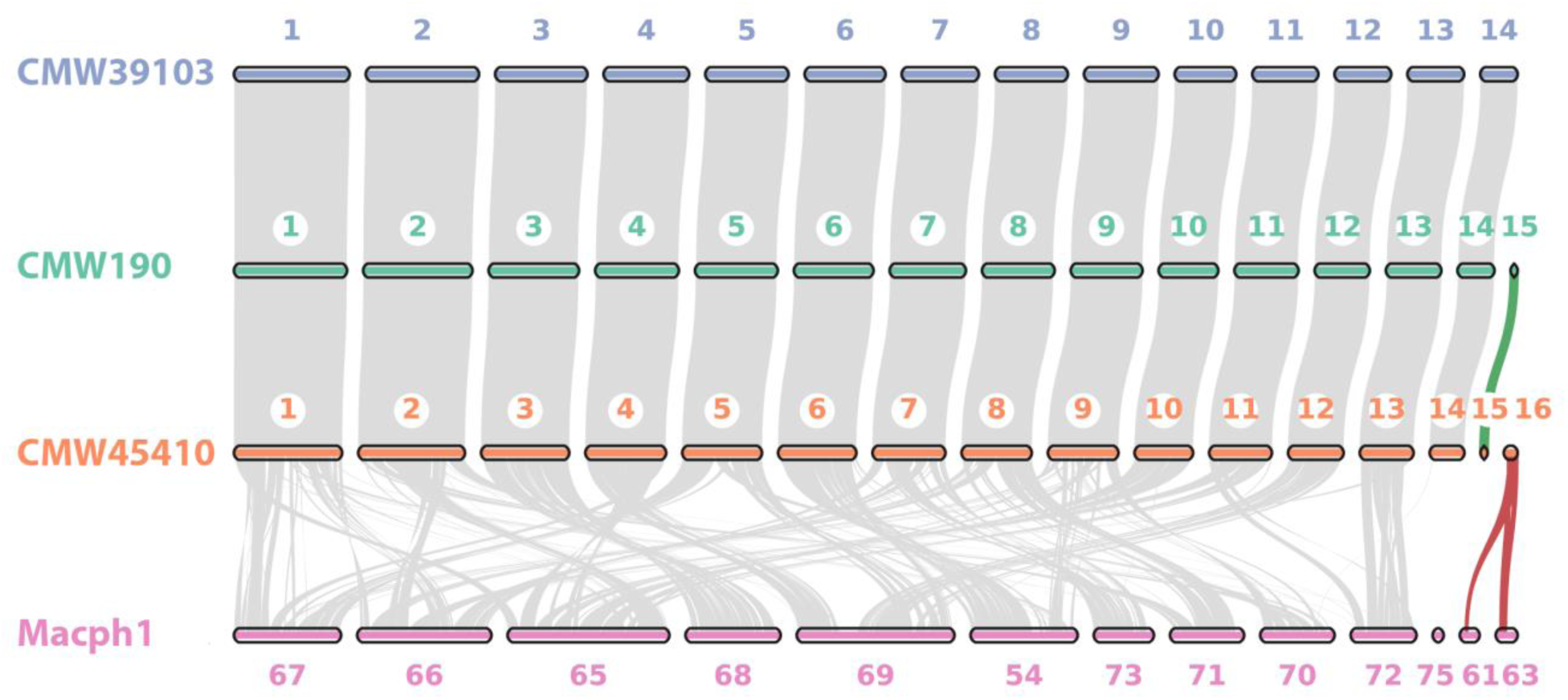
Genome-wide collinearity between three assembled *D. sapinea* genomes and assembled genome of *Macrophomina phaseolina* (GCA_020875535.1). Colinear blocks were identified and visualized using MCScanX (Wang et al. 2012). Putative chromosome numbers are assigned for *D. sapinea* genomes, and contig numbers in the original *M. phaseolina* genome were used.

### Genes on the ACs display high sequence homology to *Macrophomina phaseolina*, a distantly related species of *Botryosphaeriaceae*

Sequence homology analysis indicated that most of the proteins encoded by genes of the core chromosomes (CCs) had above 90% similarity to available sequences on the NR database, whereas most of the proteins encoded by genes on the ACs displayed much lower sequence similarity, averaging 68.4% (Figure 2B). Protein sequences encoded by genes on the CCs showed the highest similarity to other *Diplodia* sp., with *Diplodia seriata* accounting for over 75% of the top five species BLAST hits, followed by *Diplodia corticola* with over 15% (Figure 2C). This was expected, as reflected by the close evolutionary relationship between *D. sapinea*, *D. seriata* and *D. corticola* (Figure 2A). Although phylogenomic analysis indicated that *D. scrobiculata* is more closely related to *D. sapinea* than *D. corticola* and *D. seriata*, amino acid sequences for *D. scrobiculata* were not available in the public NR database and, therefore, did not appear in the BLAST results. The ACs encoded proteins exhibited the highest similarity to *Macrophomina phaseolina,* which constituted over 31% of the top five species BLAST hits for Chr 15 and over 63% of the hits for Chr 16. The next most similar species to Chr 15 was *D. corticola* with over 21% of the hits followed by *D. seriata* with over 19% of hits. For Chr 16, the species *Lasiodiplodia theobromae* showed the next highest similarity, contributing over 20% of BLAST hits, followed by *D. corticola* with over 9% of hits.

**Figure 2.**
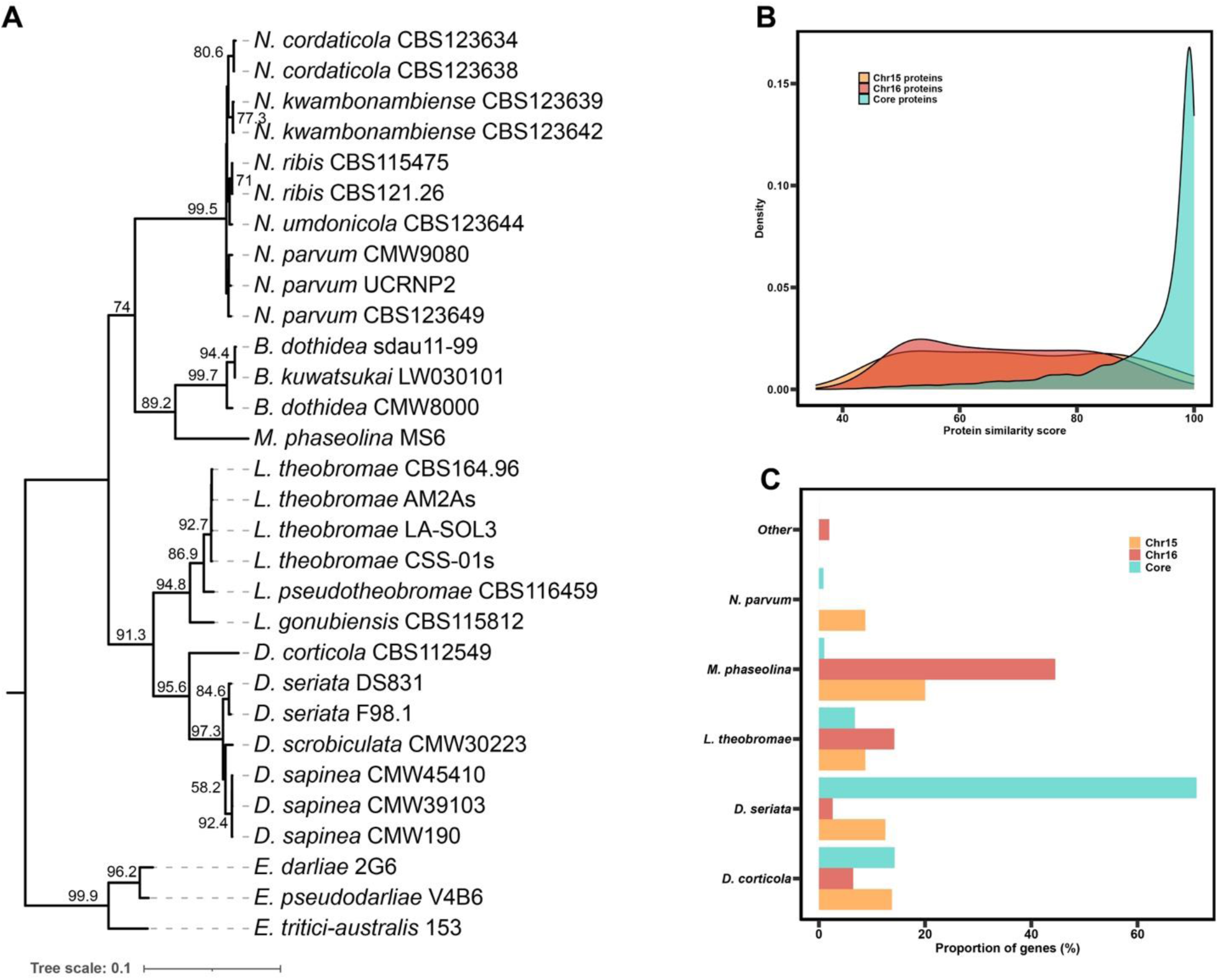
**(A)** Phylogenomic tree based on shared single copy orthologs determined by BUSCO of 30 *Botryosphaeriaceae* isolates. Gene and site concordance factors (gCF/sCF) as determined by IQ-TREE 2 are presented on the branches of the tree. **(B)** Distribution of BLAST similarity scores against NR protein database for putative amino acid sequences encoded by genes of each AC compared to those of the core chromosomes. **(C)** Proportion of proteins attributed to the top 5 species BLAST hits for putative proteins encoded by genes on each AC and the core genome.

### The ACs display evolutionary plasticity and are enriched in effectors

The genomic composition (genes, transposable elements, noncoding) of the ACs was distinct from that of the core chromosomes (Figure 3A). Both ACs had a higher abundance of TEs and lower gene densities when compared to that of the core chromosomes.

**Figure 3.**
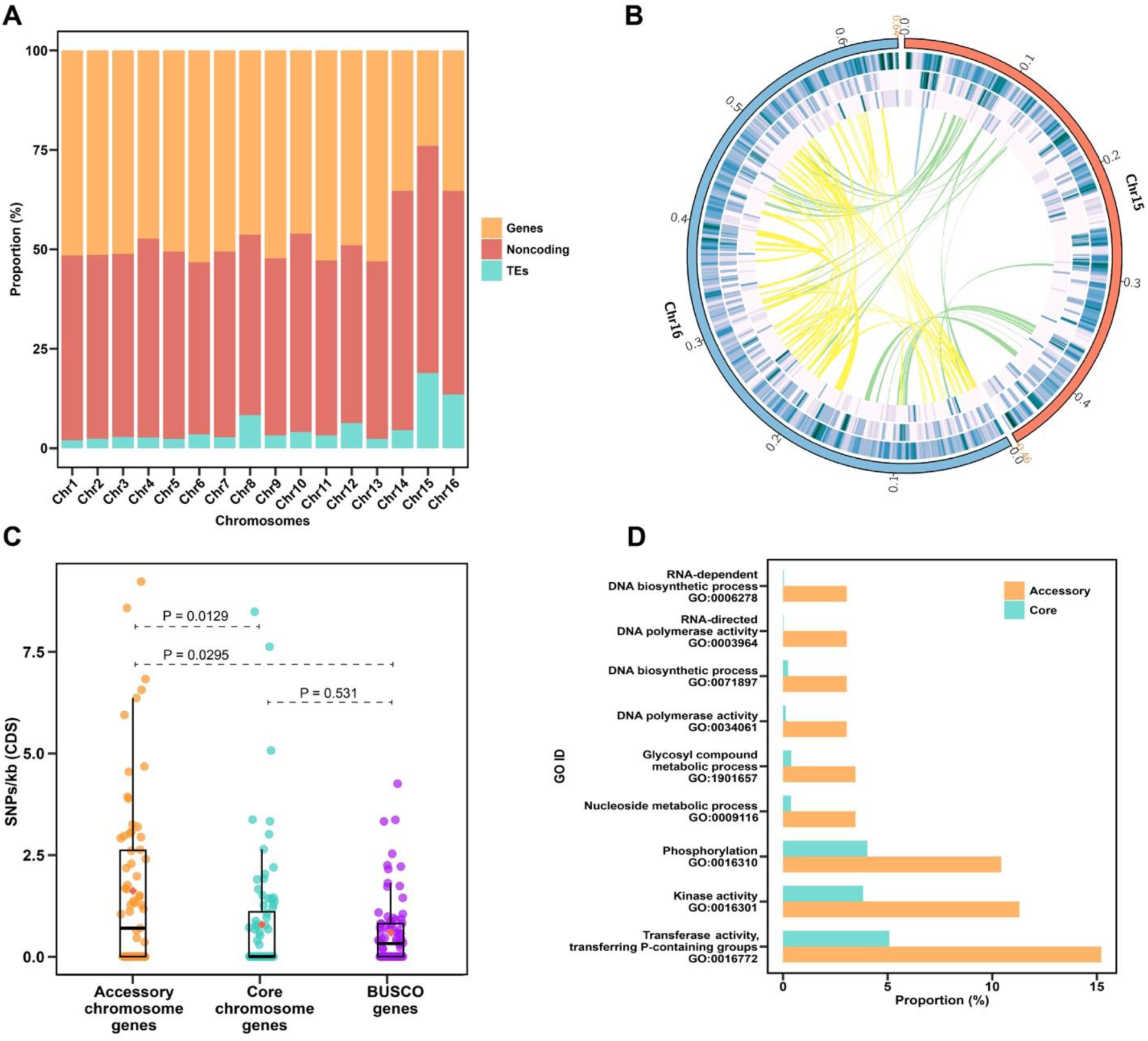
**(A)** Genomic composition (genes, noncoding and transposable elements (TEs) of the 16 pseudochromosomes in *D. sapinea* isolate CMW45410. The ACs are labelled as Chr 15 and Chr 16. **(B)** Circular visualisation of the ACs in isolate CMW45410 ACs. Tracks from outer to inner: 1: Accessory chromosomes and their lengths in Mb; 2: GC density; 3: Gene density; 3: Gene density; 4: TE density. Links show orthologous genes between the core and ACs (green = between Chr 15 and Chr 16, blue = Within Chr 15, yellow = Within Chr 16). **(C)** Number of SNPs per Kb of CDS for genes on Chr 15, a random subset of 80 genes from core chromosomes and 80 random BUSCO genes. Statistical significance was calculated using the Wilcoxon rank-sum test. **(D)** GO enrichment analysis of genes on the accessory chromosomes versus genes on the core chromosomes.

Approximately 24% of Chr 15 and 33.5% of Chr 16 were comprised of genes, while transposable elements constituted approximately 18.9% and 13.5% of these chromosomes, respectively. In contrast, all CCs consisted of at least 46% gene regions, with the exception of Chr 14, which was 35.3%. Among the CCs, Chr 8 displayed the highest proportion of TEs, accounting for 8.3% of its composition.

Sequence comparison indicated that genes on AC Chr 15 had a significantly lower level of sequence conservation when compared to CC genes (Figure 3C). Additionally, 86% of the identified SNPs in genes on the AC were RIP-like mutations (C ◊ T/G ◊ A). This is more than double compared to the RIP-like mutations identified in genes on the core chromosomes (Supp. Figure S4). Sequence variation analysis could not be conducted for Chr 16 due to it being present in only a single isolate.

Protein clustering analysis of proteins encoded by the ACs revealed that 80 putative proteins encoded by Chr 16 share homology, indicating that this chromosome has undergone multiple segmental duplications (Figure 3B). In contrast, only four putative proteins of Chr 15 shared homology with one another. When comparing between the ACs, 19 putative proteins share homology with 29 proteins of Chr 16.

A total of 1147 secreted proteins were identified in isolate CMW45410, 8 of which originated from Chr 15 and 4 from Chr 16. A further 12 unconventionally secreted proteins were identified for Chr 15 and 29 for Chr 16. Of the predicted secreted peptides, 11 were predicted as effectors for Chr 15 and 18 were predicted as effectors for Chr 16. The ACs had a higher prevalence of putative effectors, with 13.75% of genes on Chr 15 and 11.84% of genes on Chr 16 encoding putative effector proteins, compared to 8.92% of genes on the CCs encoding putative effector proteins. A fungal-RiP-like gene cluster was also predicted on Chr 16, and no biosynthetic gene cluster was detected on Chr15.

Gene ontology enrichment analysis between genes on ACs and core chromosomes showed that the ACs are enriched for genes associated with phosphorous and phosphate group utilization (Figure 3D). The most prevalent GO terms include transferase activity, kinase activity and phosphorylation. Many of the enriched GO terms for the ACs appear to be associated with TE activities, including RNA and DNA-dependent polymerase activities and biosynthetic processes.

### Chr 15 was widespread whereas Chr 16 was detected in only a single isolate

Low-coverage Illumina sequencing of six *D. sapinea* isolates (Supp. Table S1) showed that Chr 15 occurred commonly and was found in eight of the nine isolates for which genome data was generated. These isolates were from South Africa, Sweden, Italy, Chile and France. One isolate (CMW39103) from South Africa did not contain this chromosome. In contrast, the second AC was found in only a single isolate (CMW45410) originating from Sweden. Multiplex PCR assays targeting three genes on each of the ACs on a total of 40 isolates (including CMW190, CMW39103 and CMW45410) resulted in the identification of Chr 15 in 35 isolates. None of the additional isolates investigated harboured Chr 16 (Supp. Table S4). Whole genome alignment of three recently generated *D. sapinea* genomes from China (KE8364, KE8391 and KE8634) (Yu et al. 2022) with the genome of CMW45410 revealed that all three isolates possess AC Chr 15, and none have Chr 16. Isolates from the 17 countries considered for the presence of ACs showed that 16 contained the AC Chr 15. These isolates were from South Africa, Ethiopia, New Zealand, Montenegro, Serbia, Sweden, Switzerland, the USA, Brazil, Chile, Colombia, China, Netherlands, Great Britain, France and Italy. Isolates from Indonesia did not contain either of the ACs. A table summarising AC presence for all isolates is available in Supp. (Supp. Table. S7).

### The presence of ACs showed no correlation with pathogenicity

The *D. sapinea* isolates used in the inoculation trial showed significant variation in their ability to produce stem lesions, but there was no clear correlation between AC presence and lesion length. Regardless of whether ACs were present or absent, most isolates produced clear lesions longer than those of the controls. The only exception was isolate CMW29644 that produced relatively small lesions (Supp. Figure S5). The isolates without the ACs (CMW39103, CMW34220, and CMW4889) produced average lesion lengths of 68.71 mm, 67.14 mm and 57.86 mm, respectively (Supp. Table. S8). The isolates (CMW8754 and CMW34220) having AC Chr 15 produced average lesion lengths of 58.71 mm and 19 mm, respectively. The isolate (CMW 45410) with both AC Chr 15 and Chr 16, gave rise to lesions having an average length of 62 mm. None of the control inoculated plants had lesions. Re-isolations, followed by sequencing of the ITS region, confirmed the presence of *D. sapinea* on the dead trees.

## Discussion

Chromosome-level genome assemblies were generated for three isolates of *D. sapinea* and genomic data were produced for six additional isolates. No other assemblies for this species, or any other Botryosphaeriaceae, have reached the chromosome level of completeness. This level of genome completeness, contiguity, and structural accuracy allowed for the discovery of two ACs for the first time in this pathogen. Sequence analyses revealed that these ACs have unique genomic compositions compared to the core chromosomes and that they have likely been acquired horizontally. Pathogenicity assays showed no correlation between the presence of ACs and the length of lesions on inoculated saplings.

The ACs identified in this study are the first to be identified in a species of the *Botryosphaeriaceae*. Comparison of protein sequences encoded by genes of the ACs and CCs with available proteins from the NR database revealed different profiles of taxonomic association. Proteins encoded by CCs showed the highest similarity to proteins in other *Diplodia* species, while proteins encoded by the AC genes showed higher similarity to those from *M. phaseolina*, a distantly related species in the *Botryosphaeriaceae*. This suggests a potential horizontal origin for both ACs discovered in *D. sapinea*. Additionally, the unique genomic composition of the ACs compared to the CC also suggests that they have been acquired by horizontal transfer. Alternatively, these ACs could be common in members of the *Botryosphaeriaceae* and be inherited vertically. If this is true, they will be identified in other members of the family as more complete genomes are generated.

The ACs identified in this study exhibited distinct genomic characteristics compared to the CCs. For example, the ACs displayed a lower gene density, a higher proportion of TEs, and increased SNP variations. Such higher TE content is typically observed for ACs (Williams et al. 2016) and can be attributed to the fact that they do not carry essential genes, thereby minimizing the risk of TE disrupting vital gene functions (Bertazzoni et al. 2018). The presence of TEs can contribute to genome plasticity and generate further variation via RIP mutations (Lorrain et al. 2021). The segmental duplications observed in Chr16 are likely mediated by TEs. Orthology analysis indicated that half of the proteins on Chr 16 show orthology to each other as the result of these duplications. These duplicated genes can serve as sources for functional diversification (Marques et al. 2008), suggesting the adaptive evolutionary roles of ACs.

The ACs discovered in *D. sapinea* align with several other similar discoveries in plant pathogens that have recently been shown to contain ACs. ACs in plant pathogens are thought to potentially play an important role as a driving force in genome evolution, affecting important phenotypic traits such as pathogenicity, virulence, and host adaptation. Examples of this include an accessory chromosome in *Nectria haematococca* that results in increased pathogenicity to *Pisum sativum* (Han et al. 2001). A pathogenicity gene cluster that resides on this AC contains the PDA1 gene that provides resistance to pisatin, an antimicrobial produced by the host plant during infection (Han et al. 2001). Similarly, Ma et al. (2010) showed that isolates of *Fusarium oxysporum* without an AC were initially able to penetrate host cells but were soon suppressed during the biotrophic phase, suggesting that the AC contained genes coding for proteins involved in overcoming host defences post host penetration. More recently, *C. higginsianum* was shown to possess two ACs, one of which was directly associated with pathogenicity to *Arabidopsis thaliana* (Plaumann et al. 2018).

In contrast to the abovementioned studies, no association was found between the presence of ACs in *D. sapinea* and pathogenicity. Furthermore, the differences in isolate pathogenicity despite the presence or absence of ACs suggest that other genetic mechanisms unrelated to the ACs may be responsible for the observed differences in disease development. It is possible, however, that the ACs only produce a discernible effect under particular environmental conditions, host conditions or host species (as discussed in Bertazzoni et al. 2018), which were not met in the pathogenicity trial of this study. Alternatively, these ACs may not contribute directly towards pathogenicity but play a role in the responses of *D. sapinea* to other biotic or abiotic factors.

It is possible that the ACs discovered in this study play a role in the ability of *D. sapinea* to colonize its host endophytically. In *Stagonosporopsis rhizophila,* for example, Wei et al. (2024) showed that the loss of an AC changed the phenotype of this fungus from being pathogenic to endophytic. *Diplodia sapinea* is known to colonize its host without causing symptoms and to exist as a latent pathogen (Bihon et al. 2011). Given the common nature of this characteristic in *D. sapinea*, AC Chr 15 would be the most likely candidate to be involved in the process because it is present in most isolates. The AC Chr 15 also has a higher proportion of effectors compared to both the CCs and Chr 16, and these effectors play a role in invading host immunity or manipulating host responses (Li et al. 2024).

The two ACs found in isolates of *D. sapinea* were not distributed equally amongst the isolates investigated. Chr 15 occurs at a high frequency across multiple populations that span two countries in Africa, eight cs in Europe, one in Asia, three in South America, one North American country and one in Oceania. In contrast, Chr 16 was found only in a single isolate of *D. sapinea* isolate from Sweden. One explanation for these different patterns of association is that the acquisition of Chr 16 may have been a recent event, and as a result, this AC has not become established within all *D. sapinea* populations. Alternatively, this AC may only provide an advantage under particular conditions (Bertazzoni et al. 2018) and is only maintained at high frequency in certain populations of *D. sapinea*.

The provision of a chromosome-level assembly for *D. sapinea* in this study represents a valuable resource providing for new avenues of investigation on t this pathogen in general and contributes to its potential for the development of an experimental system for this latent tree pathogen. For example, Oostlander et al. (2023, 2024) developed sporulation, infection and transformation tools for the study of infection and host-pathogen interaction that can now be exploited more precisely to study of gene function in this process. In addition, the discovery of ACs in *D. sapinea* provides intriguing new opportunities to gain insights into the biology and evolution of this pathogen through reverse genetics or through the curing of isolates from one or more of these ACs. Collectively, these new avenues of study should answer the many questions relating to the latent and pathogenic lifestyles of *D. sapinea*.

## Data Availability Statement

The genome assemblies of three *D. sapinea* strains have been deposited in NCBI GenBank database and accession numbers are PENDING. The short and long read sequence data have been deposited in NCBI Sequence Read Archive (SRA) and the accession numbers are provided in Table S1. The pipelines and scripts used for data analyses are available on GitHub: https://github.com/PLockeS/Dsapinea_accessory_chromosomes. Genome assemblies, annotations and variant call file have been deposited in FigShare: https://figshare.com/s/24a621f6b1e3d338d31f.

## Acknowledgements

Cultures of *Diplodia sapinea* used in this study originated from collaborations with colleagues from many different parts of the world and for which we are most grateful. We also recognise the staff of the culture collection (CMW) of the Forestry and Agricultural Biotechnology Institute (FABI) at the University of Pretoria for maintaining these cultures and making them available for this study.

## Funding

This research was funded by members of the Tree Protection Co-operative Program (TPCP) and the DSI-NRF SARChI Chair in Fungal Genomics. Benoit Laurent was funded by COTE Cluster of Excellence Grant Number ANR-10-LABX-45.

## Supplementary material

**Table S1.** Isolates of *D. sapinea* used for genome sequencing and analyses.

| Isolate | Host | Country of isolation | Illumina accession | Nanopore accession |
| --- | --- | --- | --- | --- |
| CMW39103 | *Pinus patula* | South Africa | SRR32830896 | SRR32830895 |
| CMW190 | *Pinus banksiana* | South Africa | SRR32830892 | SRR32830891 |
| CMW45410 | *Pinus nigra* | Sweden | SRR32830890 | SRR32830889 |
| CBS109727 | *Pinus radiata* | South Africa | SRR32830888 | N/A |
| CBS119938 | *Pinus radiata* | Italy | SRR32830887 | N/A |
| CBS120833 | *Prunus persica* | South Africa | SRR32830886 | N/A |
| CBS623.74 | *Pinus radiata* | Chile | SRR32830885 | N/A |
| 1684n<sup>e</sup>6-1 | *Pinus pinaster* | France | SRR32830894 | N/A |
| Pier4 | *Pinus nigra* | France | SRR32830893 | N/A |

**Table S2.**
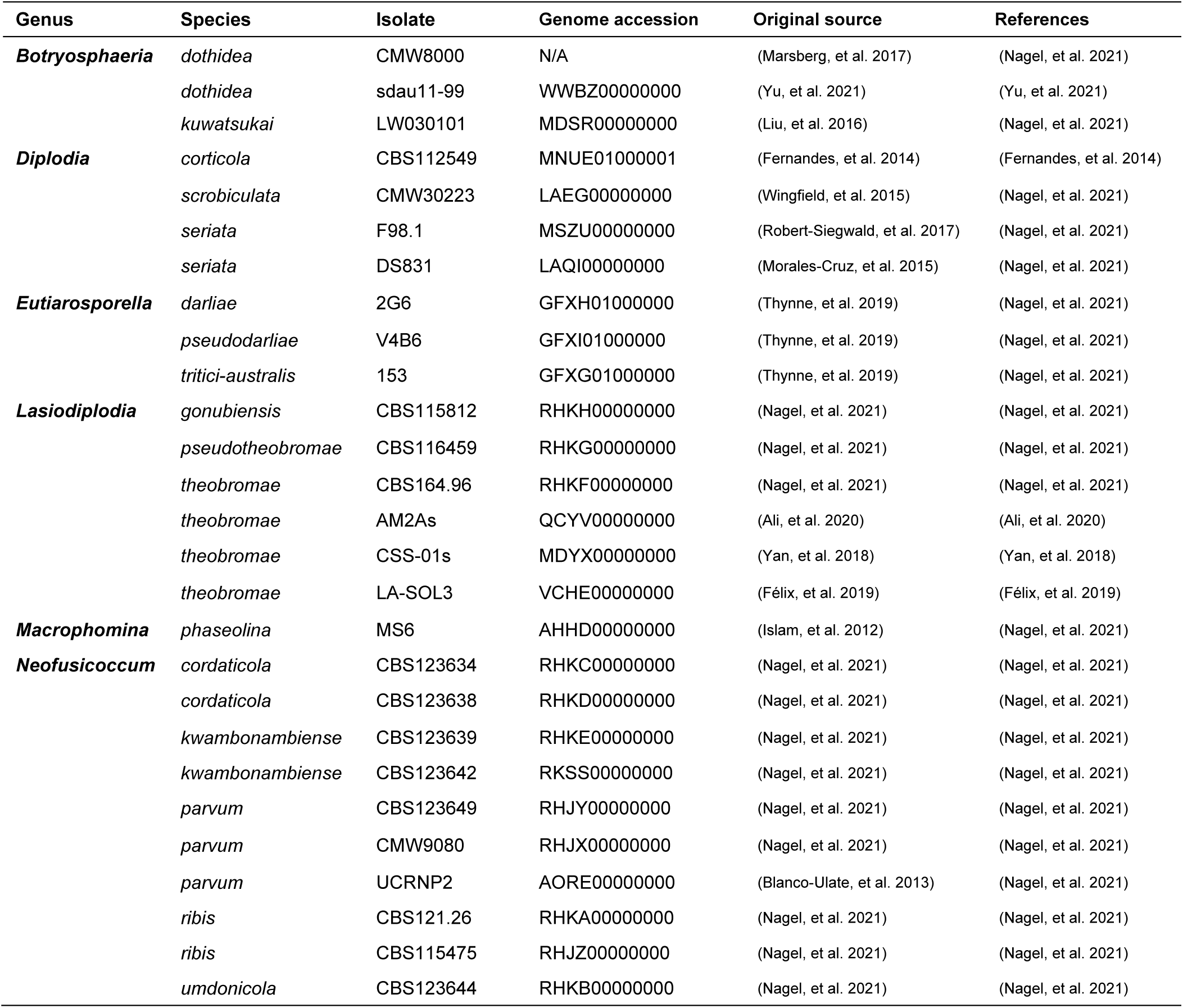
Species of *Botryosphaeriaceae* used in protein clustering and phylogenomic analysis.

**Table S3.**
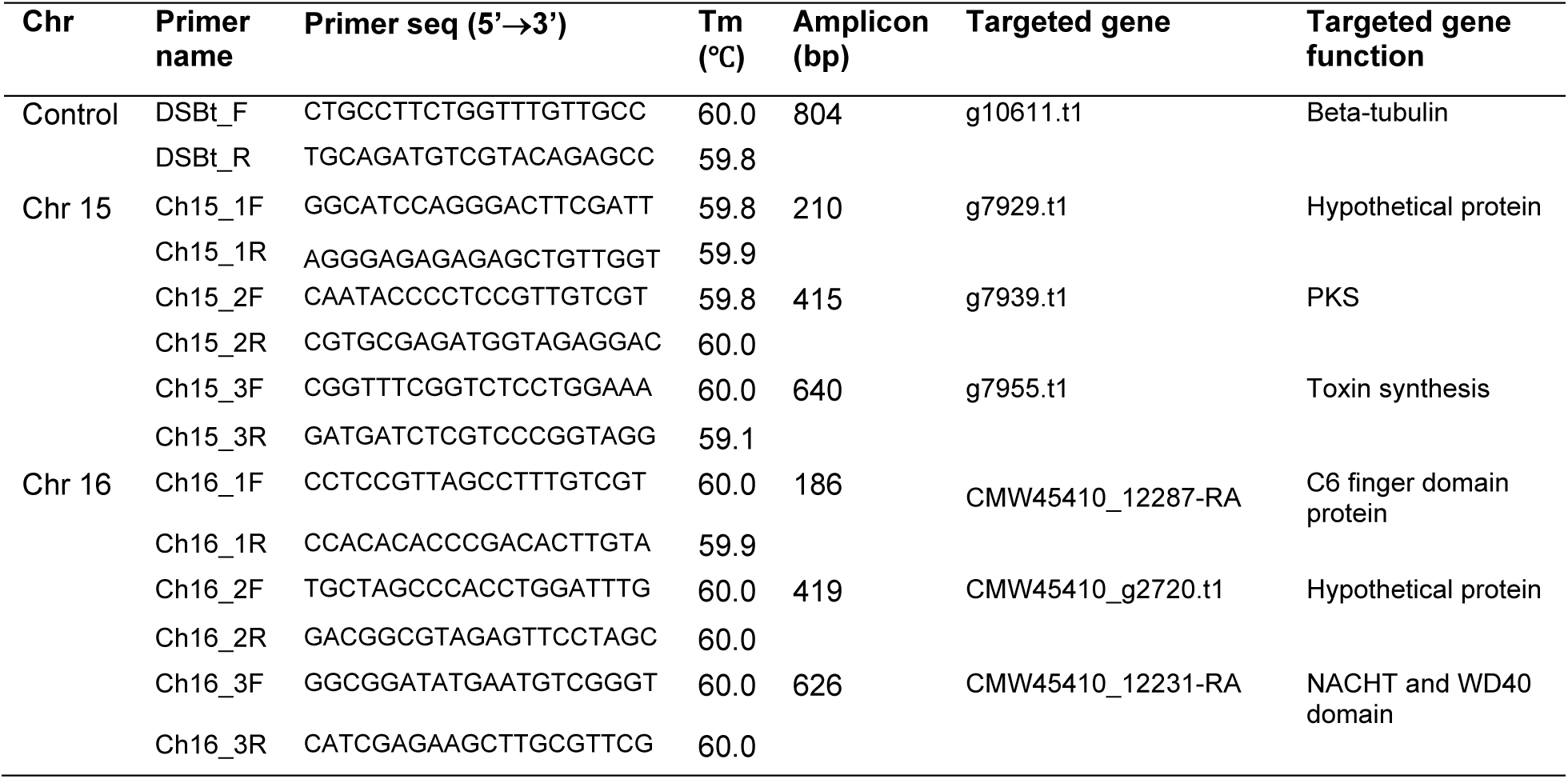
Primer design for detection of ACs in *D. sapinea*.

**Table S4.** Results of PCR assays to identify dispensable chromosomes in *D. sapinea* isolates from various countries around the world.

| Isolate (CMW) | Chr 15 | Chr 16 | Country |
| --- | --- | --- | --- |
| 39103 | - | - | South Africa |
| 190 | + | - | South Africa |
| 45410 | + | + | Sweden |
| 29644 | + | - | Ethiopia |
| 4237 | + | - | South Africa |
| 29133 | + | - | South Africa |
| 29309 | + | - | South Africa |
| 29311 | + | - | South Africa |
| 31572 | + | - | South Africa |
| 31649 | + | - | South Africa |
| 31655 | + | - | South Africa |
| 32008 | + | - | South Africa |
| 34302 | + | - | South Africa |
| 24705 | + | - | China |
| 24707 | + | - | China |
| 24708 | + | - | China |
| 4889 | - | - | Indonesia |
| 32287 | - | - | Indonesia |
| 32361 | - | - | Indonesia |
| 34220 | - | - | Indonesia |
| 8750 | + | - | Great Britain |
| 33544 | + | - | Great Britain |
| 39331 | + | - | Montenegro* |
| 39333 | + | - | Montenegro |
| 39337 | + | - | Montenegro |
| 39341 | + | - | Montenegro |
| 8749 | + | - | Netherlands |
| 33543 | + | - | Netherlands |
| 39329 | + | - | Serbia |
| 39338 | + | - | Serbia |
| 39342 | + | - | Serbia |

| **Isolate (CMW)** | **Chr 15** | **Chr 16** | **Country** |
| --- | --- | --- | --- |
| 32277 | + | - | Switzerland* |
| 8754 | + | - | USA |
| 31975 | + | - | Brazil |
| 14971 | + | - | Chile |
| 14974 | + | - | Chile |
| 5850 | + | - | Colombia |
| 32112 | + | - | Colombia |
| 32425 | + | - | New Zealand |
| 32431 | + | - | New Zealand |

**Table S5.** Summary of *D. sapinea* isolates used in pathogenicity trials.

| Isolate (CMW) | Origin | Host | DCs |
| --- | --- | --- | --- |
| 39103 | South Africa | *P. patula* | - |
| 4889 | Indonesia | *P. patula* | - |
| 34220 | Indonesia | *P. patula* | - |
| 29644 | Ethiopia | *P. patula* | Chr 15 |
| 8754 | USA | *Pseudotsuga menziessi* | Chr 15 |
| 45410 | Sweden | *P. nigra* | Chr 15, Chr 16 |

**Table S6.** Statistics of genome assemblies produced by Canu and Flye for each *D. sapinea* isolate.

| Isolate | Assembler | Contigs | Size (Mb) | N50 | L50 | Mis-joins | Splits |
| --- | --- | --- | --- | --- | --- | --- | --- |
| CMW39103 | Canu | 14 | 36.88 | 2859033 | 6 | 0 | 0 |
|  | Flye | 13 | 36.7 | 2852626 | 6 | 1 | 0 |
| CMW190 | Canu | 16 | 37.98 | 2831583 | 6 | 0 | 1 |
|  | Flye | 13 | 37.27 | 3188804 | 5 | 2 | 0 |
| CMW45410 | Canu | 18 | 38.07 | 2676158 | 7 | 0 | 2 |
|  | Flye | 15 | 37.93 | 2861173 | 5 | 1 | 0 |

**Table S7.**
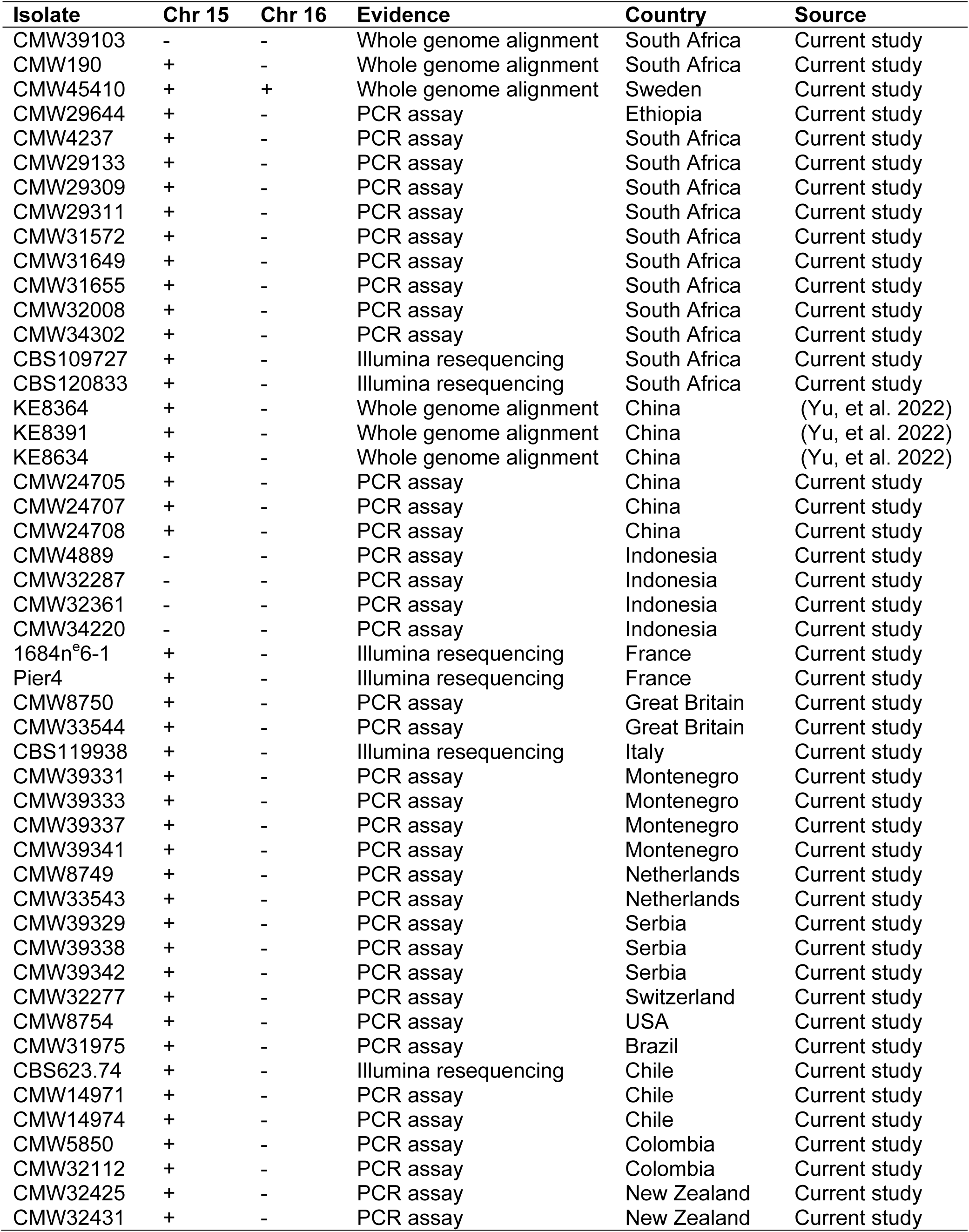
Summary of all isolates used in this study and whether the DCs are present or absent.

**Table S8.** Lesion lengths measured from pathogenicity trial.

| Isolate | ACs | Lesion lengths<br>(mm) | Average lesion length<br>(mm) |
| --- | --- | --- | --- |
| CMW39103 | None | 53 | 68.71 |
|  |  | 66 |  |
|  |  | 68 |  |
|  |  | 71 |  |
|  |  | 72 |  |
|  |  | 74 |  |
|  |  | 77 |  |
| CMW4889 | None | 7 | 57.86 |
|  |  | 13 |  |
|  |  | 57 |  |
|  |  | 65 |  |
|  |  | 78 |  |
|  |  | 88 |  |
|  |  | 97 |  |
| CMW34220 | None | 32 | 67.14 |
|  |  | 49 |  |
|  |  | 63 |  |
|  |  | 71 |  |
|  |  | 76 |  |
|  |  | 88 |  |
|  |  | 91 |  |
| CMW8754 | Chr 15 | 42 | 58.71 |
|  |  | 43 |  |
|  |  | 45 |  |
|  |  | 48 |  |
|  |  | 62 |  |
|  |  | 79 |  |
|  |  | 92 |  |
| CMW29644 | Chr 15 | 7 | 19 |
|  |  | 8 |  |
|  |  | 11 |  |
|  |  | 12 |  |
|  |  | 24 |  |
|  |  | 28 |  |
|  |  | 43 |  |
| CMW45410 | Chr 15, Chr 16 | 14 | 62 |
|  |  | 43 |  |
|  |  | 49 |  |
|  |  | 52 |  |
|  |  | 87 |  |
|  |  | 91 |  |
|  |  | 98 |  |

**Figure S1.**
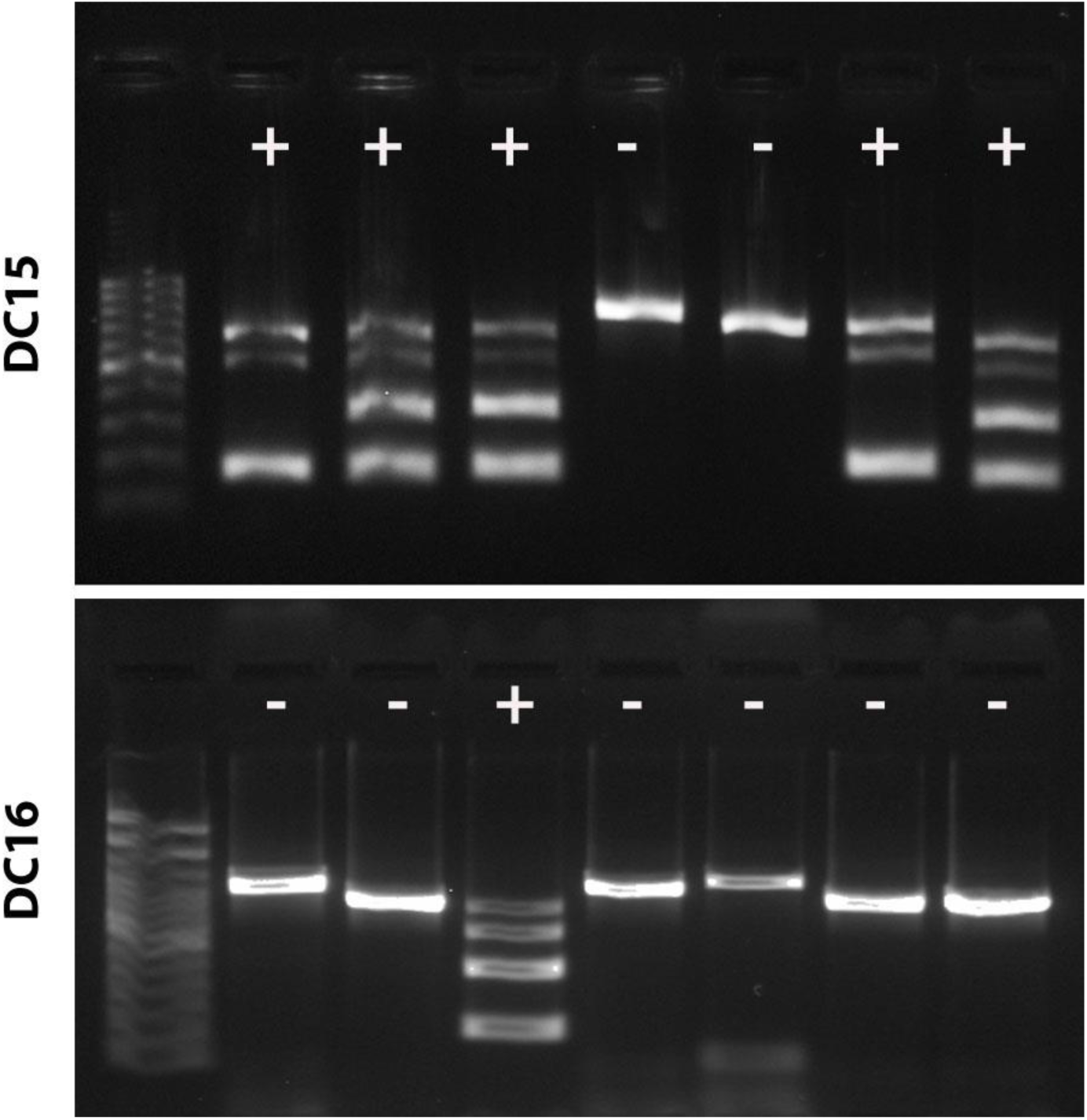
Gel electrophoresis image showing multiplex PCR assays for the detection of each AC. A single amplicon shows samples without the AC, and multiple amplicons indicate the presence of the AC.

**Figure S2.**
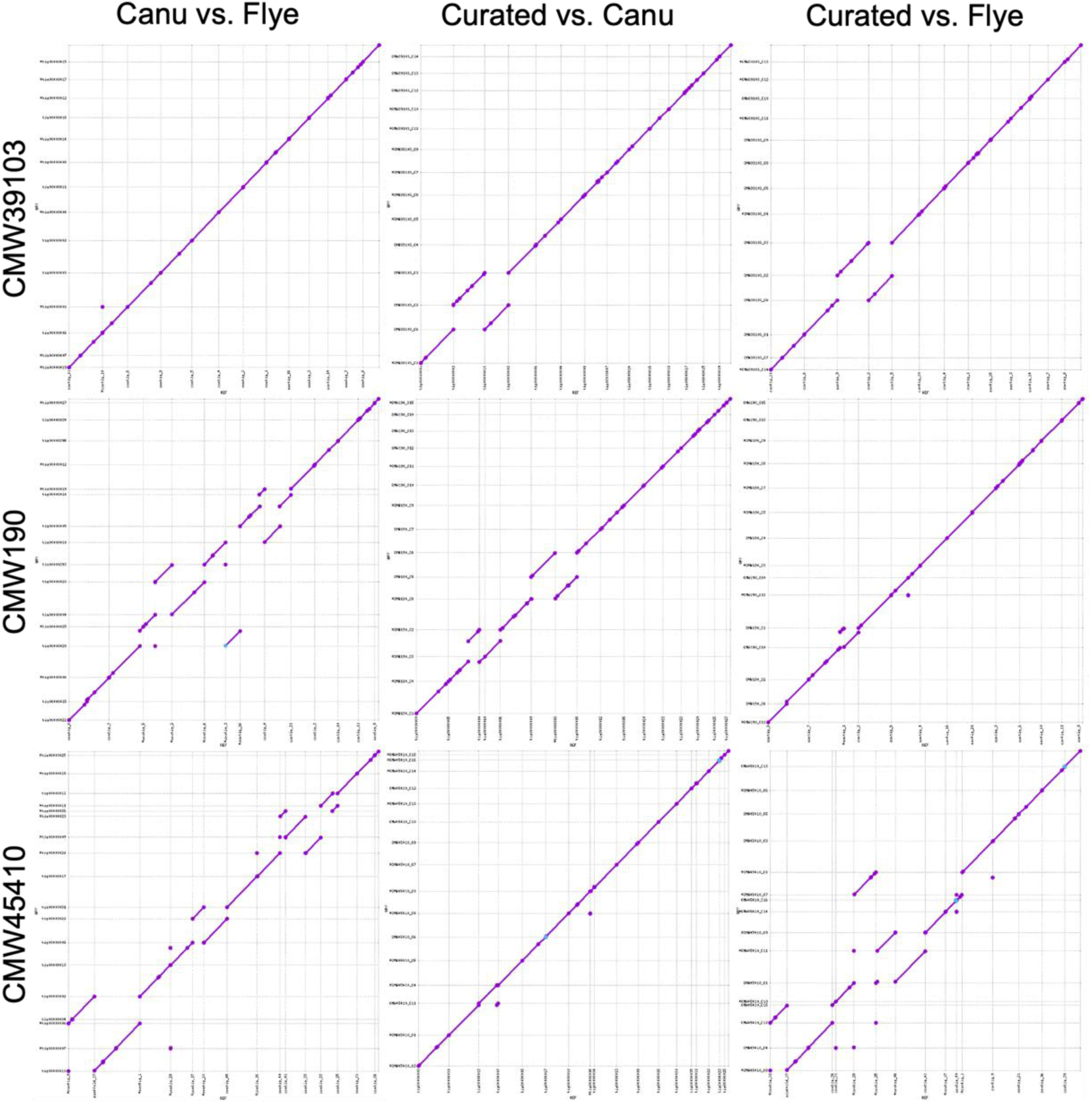
Whole genome alignments generated using MUMmer of the initial assemblies and the final curated assemblies.

**Figure S3.**
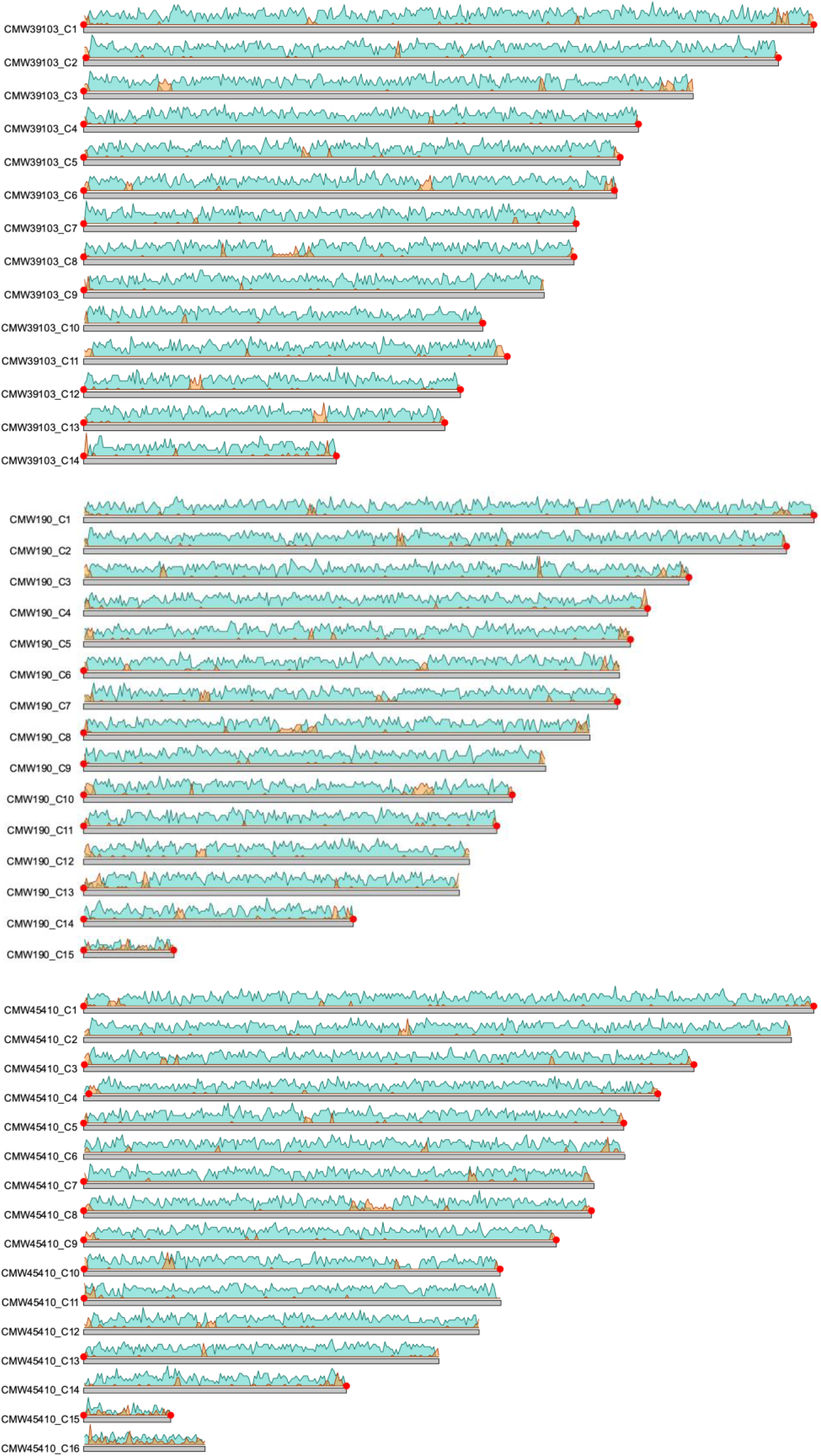
Karyoplots of the genome assemblies generated in this study for CMW39103, CMW190 and CMW45410 using karyoploteR v1.28.0 (Gel and Serra 2017). Grey bars represent the assembled contigs, blue density plots show gene density, and orange density plots show TE density. Red dots indicate an identified telomeric region.

**Figure S4.**
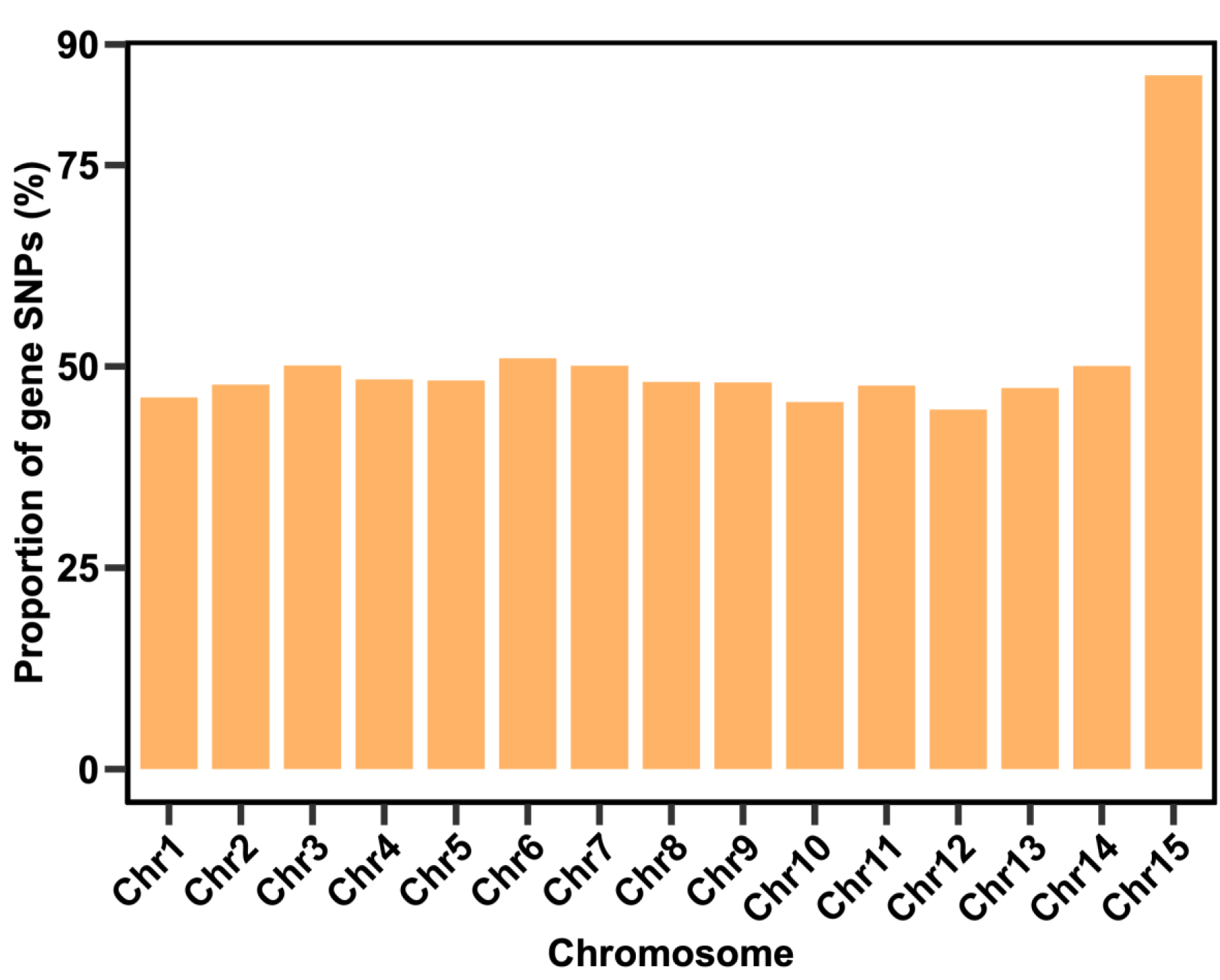
Bar graph comparing the proportion of RIP-like SNPs in gene regions between each chromosome of CMW45410.

**Figure S5.**
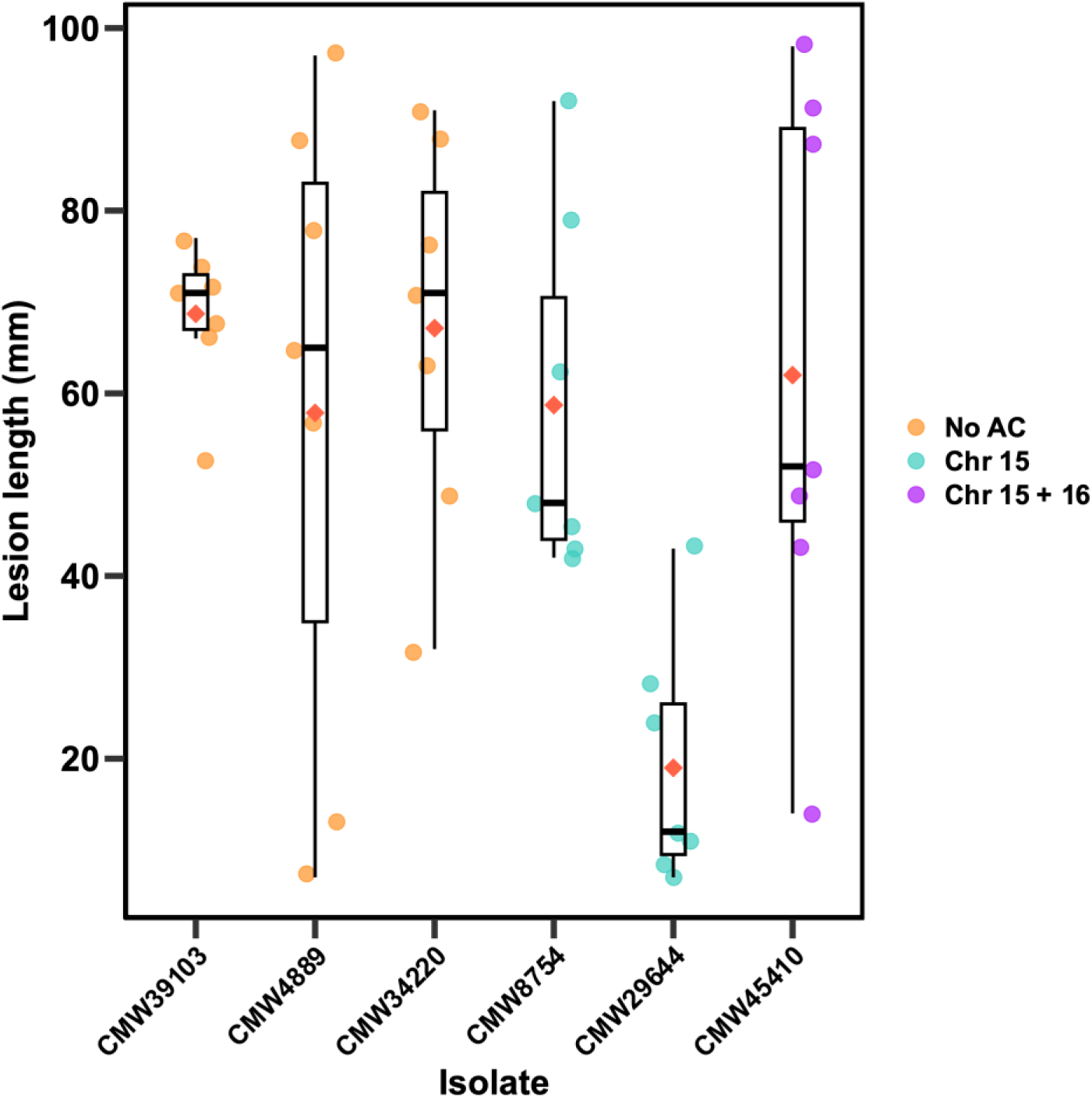
Lesion measurements produced by *D. sapinea* isolates used in the pathogenicity trial.

